# Notch3 deletion regulates HIV-1 gene expression and systemic inflammation to ameliorate chronic kidney disease

**DOI:** 10.1101/2023.09.12.557484

**Authors:** Mackenzie Thornton, Nicole Sommer, Mercedes McGonigle, Anil Kumar Ram, Sireesha Yerrathota, Henrietta Ehirim, Aakriti Chaturvedi, Johnny Dinh Phan, Anubhav Chakraborty, Praveen V Chakravarthi, Sumedha Gunewardena, Mudit Tyagi, Jaya Talreja, Tao Wang, Pravin Singhal, Pamela V Tran, Timothy A Fields, Patricio E Ray, Navneet K Dhillon, Madhulika Sharma

**Affiliations:** Departments of Internal Medicine, University of Kansas Medical Center, Kansas City, KS; Departments of Cell Biology and Physiology, University of Kansas Medical Center, Kansas City, KS; Departments of Pathology and Laboratory Medicine, University of Kansas Medical Center, Kansas City, KS; The Jared Grantham Kidney Institute, University of Kansas Medical Center, Kansas City, KS; Department of Medicine, Center for Translational Medicine, Thomas Jefferson University, Philadelphia, PA; Division of Pulmonary, Critical Care and Sleep Medicine, Wayne State University School of Medicine and Detroit Medical Center, Detroit, MI; Department of Biology, Medicine and Health, The University of Manchester, UK; Institute of Molecular Medicine, Feinstein Institute for Medical Research, Zucker School of Medicine at Hofstra-Northwell, New York, NY; Child Health Research Center and Department of Pediatrics, University of Virginia School of Medicine, Charlottesville, VA

## Abstract

Antiretroviral therapy (ART) has decreased HIV-1 associated morbidity. However, despite ART, immune cells remain latently infected and slowly release viral proteins, leading to chronic inflammation and HIV-1 associated comorbidities. New strategies are needed to target viral proteins and inflammation. We found activation of Notch3 in several renal cells of the HIV-1 mouse model (HIV-Tg26) and in patients with HIV associated Nephropathy. We hypothesized that targeting Notch3 activation constitutes an effective therapy for HIV-related chronic kidney diseases (HIV-CKD). We generated HIV-Tg26 mice with Notch3 knocked out (Tg-N3KO). Compared to HIV-Tg26 mice at 3 months, HIV-Tg-N3KO mice showed a marked reduction in renal injury, skin lesions and mortality rate. Bulk RNA sequencing revealed that N3KO not only reduced renal infiltrating cells but significantly reduced the expression of HIV genes. Moreover, Notch3 activated the HIV-promoter and induction of HIV-1 resulted in increased Notch3 activation indicating a feedback mechanism. Further, bone marrow derived macrophages (BMDMs) from HIV-Tg26 mice showed activation of Notch3 indicating systemic effects. Consistent with that, systemic levels of TNF-α, MCP-1 and other inflammatory chemokines and cytokines were reduced in Tg-N3KO mice. Thus, Notch3 inhibition/deletion has a dual therapeutic effect in HIV-CKD and may extend to other HIV-related pathologies.

## Introduction

Anti-retroviral therapy (ART) has decreased the incidence of HIV-1 related pathologies and prolonged the lives of people living with HIV-1 (PLWH). However, persistent low viremia exists in PLWH despite ART, which results in continuous immune activation and inflammation^1-4^. These events ultimately result in chronic diseases including chronic kidney disease (CKD)^4,5^. Despite this, the guidelines for preventing organ damage remain the same and do not target unique molecular pathways related to disease progression^6^. We have previously reported that renal Notch signaling is activated in patients with HIV-1 associated nephropathy (HIVAN) and in non-replicating HIV-1 transgenic rodent models^7,8^.

Notch signaling is important for cell fate decisions in development, homeostasis, and disease. Notch signaling is activated when a Notch receptor (Notch1,2,3 or 4) binds to a Notch ligand (Jagged or Delta) and initiates a series of proteolytic cleavages. The final cleavage is mediated by gamma secretase, which results in release of the Notch intracellular (NIC) domain and its translocation into the nucleus. In the nucleus, NIC binds to RBP-JK transcription factor and converts it into a transcriptional activator of *Hes* and *Hey* family genes^9-12^. Notch signaling is essential for nephrogenesis, but its suppression is necessary for terminal differentiation^13–19^. Notch overexpression has been reported in glomerular diseases including HIVAN^19–23^. However, most studies have focused on targeting the gamma secretase complex which also targets other signaling pathways including Wnt and mTOR ^24,25^, and thus has yet not been successful in clinical trials. An approach to identify and target a specific disease-related Notch member holds promise. The HIV transgenic mouse (HIV-Tg26) and rat models are considered clinically relevant pre-clinical models to study co-morbidities in PLWH on ART since these models, express HIV proteins under the LTR promoter (except *gag* and *pol*) but lack viral replication. These small animals exhibit disease that mimics ART-controlled HIV-infected patients from the continuous stress of viral proteins ^26–31^. Targeted deletion of Notch1 or Notch2 is embryonic lethal, whereas mice with global deletion of Notch3 or Notch4 develop normally^32–35^. Thus, targeting Notch3 or 4 appears to be safe. In our previous studies global targeting of the Notch4 in the HIV-Tg26 mouse model led to decreased kidney injury and increased survival^36^. In the present study, we took a detailed approach to study if modulation of Notch3 axis alone holds promise.

We evaluated the effects of *Notch3* knockout (N3KO) in the HIV-Tg26 mice. Our data show that N3KO extends the life span of HIV-Tg26 mice and alleviates renal pathology and function, better than *Notch4* deletion. N3KO led to a marked reduction in the renal infiltrating cells and decreased the expression of HIV-1 genes in kidneys. Bone marrow derived macrophages (BMDMs) from HIV-Tg26 mice had activated Notch3. Finally, N3KO led to a systemic reduction of inflammatory markers, which may be an underlying mechanism of improved disease, skin lesions and prolonged life expectancy. Thus, Notch3 inhibition offers a dual-protective mechanism in HIV-related inflammation.

## Results

### Notch 3 is activated in HIV-1 human and mouse kidneys

The renal expression of Notch3 was evaluated in HIV-Tg26 mice and compared to that of age matched wildtype (WT) mice. Notch3 is normally expressed in the vascular smooth muscle cells and was seen expressed in both WT and HIV-Tg26 renal sections (asterisks, Fig:1A, B). Compared to WT sections, Notch3 was brightly labeled in the parietal epithelial cells lining the Bowmans capsule and in glomerular cells (arrow, Fig:1B) of HIV-Tg26 kidney sections. In tubular regions, Notch3 was elevated in tubular epithelial and interstitial cells (arrow, Fig:1C, D). Quantitative analysis revealed a significant increase of Notch3 in both glomerular and tubular regions of HIV-Tg26 mice as opposed to WT mice (Fig:1E, F). Further, in patients diagnosed with HIVAN, we found that Notch3 was expressed and elevated in glomerular, tubular and interstitial cells (arrows, Fig: 1G, H). Quantification of renal Notch3 staining showed a significant increase in HIVAN patients compared to biopsies from normal human kidneys (NHK) (fig:1I).

**Fig. 1.**
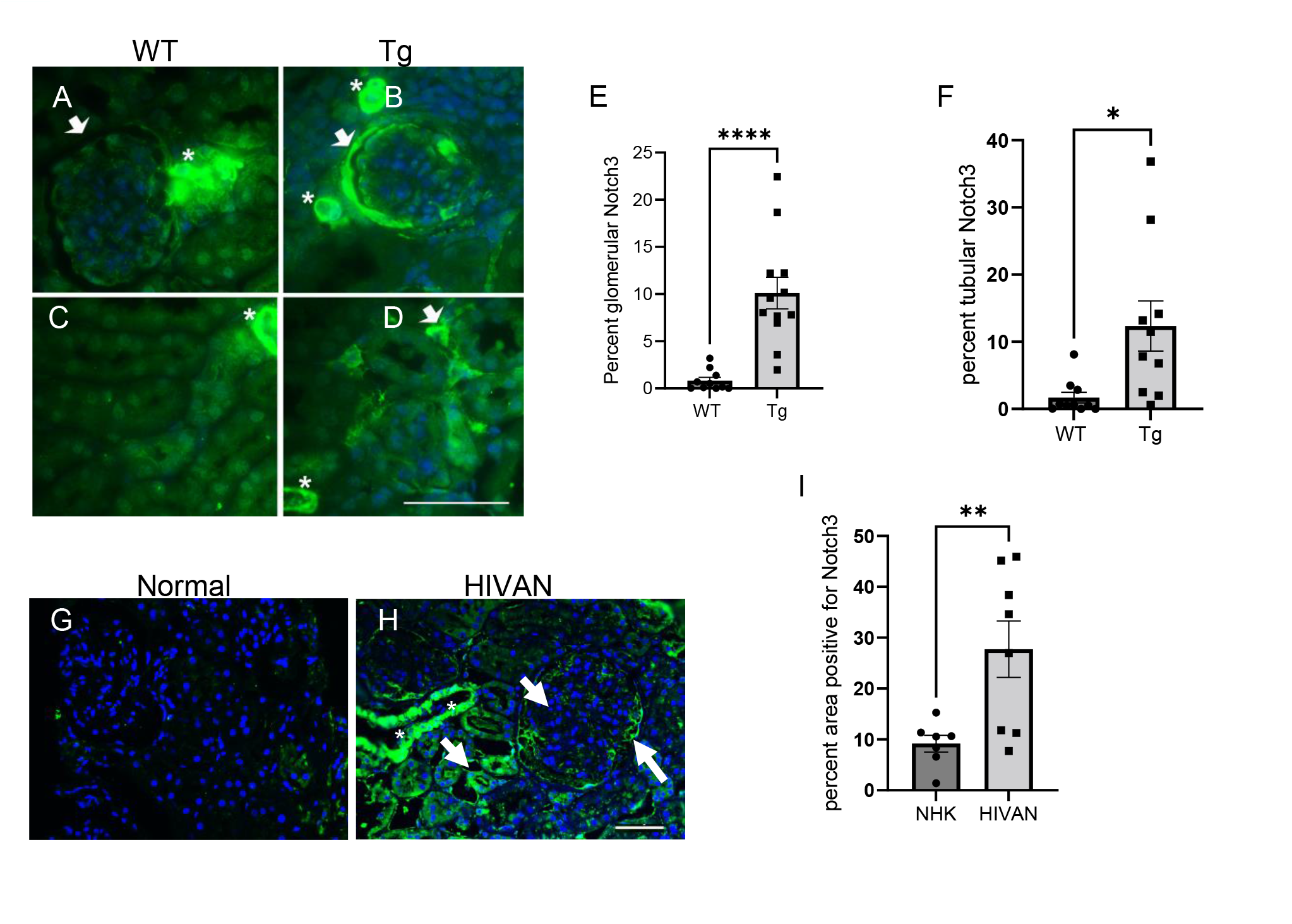
Activation of Notch3 in HIV-1 associated Nephropathy. (A-B) immunolabeling was performed for presence of Notch3 in renal paraffin sections. Notch3 (green) labeling in kidney sections of 3 months old wildtype (A and C) and Tg26 mice (B and D). Arrows indicate glomerular and tubular interstitial cells highly positive for Notch3 expression. Asterisks indicate blood vessels where Notch3 is normally expressed. (E-F) Quantification of Notch3 expression (intensity, green) as assessed using image J. (G and H) Normal versus HIVAN patient kidney biopsy (representative from n=3 in each group), nuclear expression indicating activation, arrows. (I) Quantification of Notch3 expression (intensity, green) as assessed using image J. Unpaired Student’s t-test was used, and data represented as percent area positive for green labeling (\**P*<0.05,\*\**P*<0.01, \*\*\**P*<0.001). (Scale bar: 50µm).

### Notch3 inhibition improves renal function and increases life span of the Tg26 mice

Notch3 expression was not restricted to one cell type, thus we generated Tg26 mice with global deletion of *Notch3* (HIV-Tg-N3KO) to determine *in vivo* downstream function of Notch3 activation. HIV-Tg-N3KO mice were born in the normal Mendelian fashion and were active and fertile. By 6 months of age about 10% of HIV-Tg26 mice (n=28) survived whereas 75% of HIV-Tg-N3KO mice (n=26) survived (Fig: 2A). HIV-Tg26 mice present with skin papillomata and ulcers by the age of 3 months; these abnormalities were strikingly low in HIV-Tg-N3KO mice and quantification of these showed a significant decrease in both the number and size of skin papillomas/ulcers (Fig: 2B,C, D). HIV-Tg-N3KO mice had a significant improvement in proteinuria as assessed on an SDS page gel and urinary albumin and protein creatinine ratio was drastically low in HIV-Tg-N3KO mice when compared to HIV-Tg26 mice (Fig: 2E and F). Further blood urea nitrogen levels (BUN) in HIV-Tg-N3KO mice decreased compared to the HIV-Tg26 mice showing an overall improvement in renal function (Fig. 2G).

**Fig. 2.**
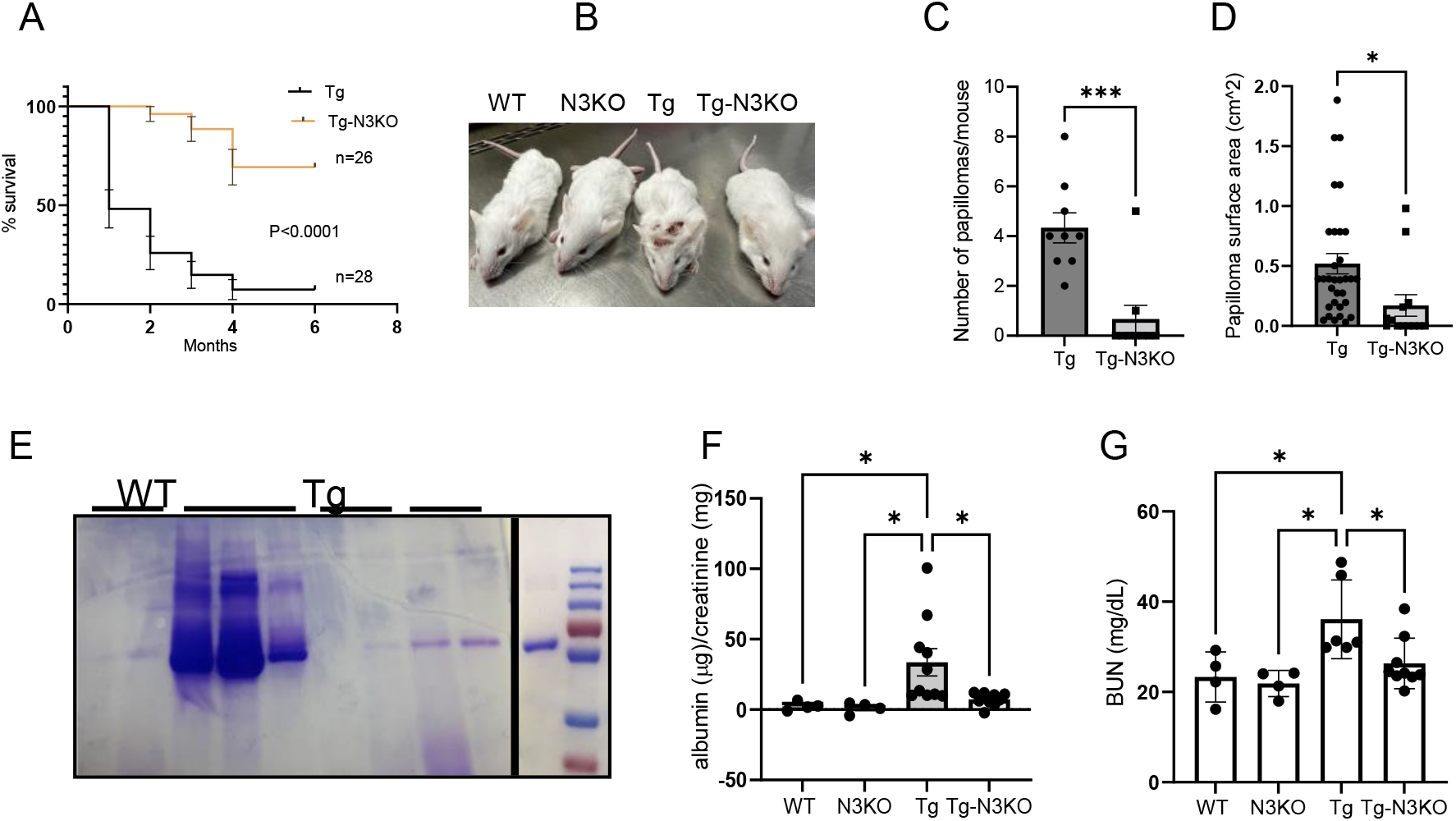
Notch deletion improves disease progression and lifespan in Tg26 mice. (A) Kaplan Meier curve showing 6 months mortality rate in Tg26 (n=26) and Tg-N3KO (n=28) mice. (B) Phenotypic appearance of WT, N3KO, Tg26 and Tg-N3KO mice, note skin papillomata on the forehead of Tg26 mouse whereas Tg-N3KO mouse appears normal. (C,D) skin lesions/mouse were quantified and area per lesion was measured and expressed as surface area per cm^2^. (E) Urine was collected in metabolic cages overnight from WT, N3KO, Tg26 and Tg-N3KO mice at 3 months of age before euthanasia. Proteinuria was assessed in 2μl urine by SDS-PAGE electrophoresis followed by Coomassie staining of the gels. “BSA (Bovine serum albumin) was used as a positive control. (F) Albumin and creatinine ratio in urine was measured using ELISA (n=4, WT) (n=4, N3KO) (n=10, Tg26) (n=10,Tg-N3KO) and expressed as μg albumin/mg creatinine. Renal function was also assessed in serum using the blood urea nitrogen (BUN) assay kit and results were expressed as BUN mg/dL (n=4, WT) (n=4, N3KO) (n=6, Tg26) (n=7,Tg-N3KO (E) (\**P*<0.05) (\*\*\**P*<0.001).

### Notch 3 deletion ameliorates renal injury in HIV-Tg26 mice

To detect histological changes in kidneys conferred by N3KO, tubulointerstitial injury, glomerular injury, and infiltrating immune cells were quantified. The N3KO mice did not show any phenotypic or histological abnormalities and were comparable to WT mice. In contrast, HIV-Tg26 mice showed tubular injury, which was reduced in HIV-Tg26-N3KO mice, though these changes did not reach statistical significance (Fig. 3A, D). Glomerular injury in HIV-Tg-N3KO mice was significantly reduced (Fig. 3B, E). The immune cell infiltration was also drastically reduced in HIV-Tg-N3KO mice compared to the HIV-Tg26 mice (Fig: 3C, F). In addition, fibrotic lesions as assessed by Mason trichrome staining were highly positive in HIV-Tg26 kidneys compared to HIV-Tg-N3KO kidneys. As expected, kidneys from both WT and N3KO mice showed no fibrotic lesions (Supplementary figure S1).

**Fig. 3.**
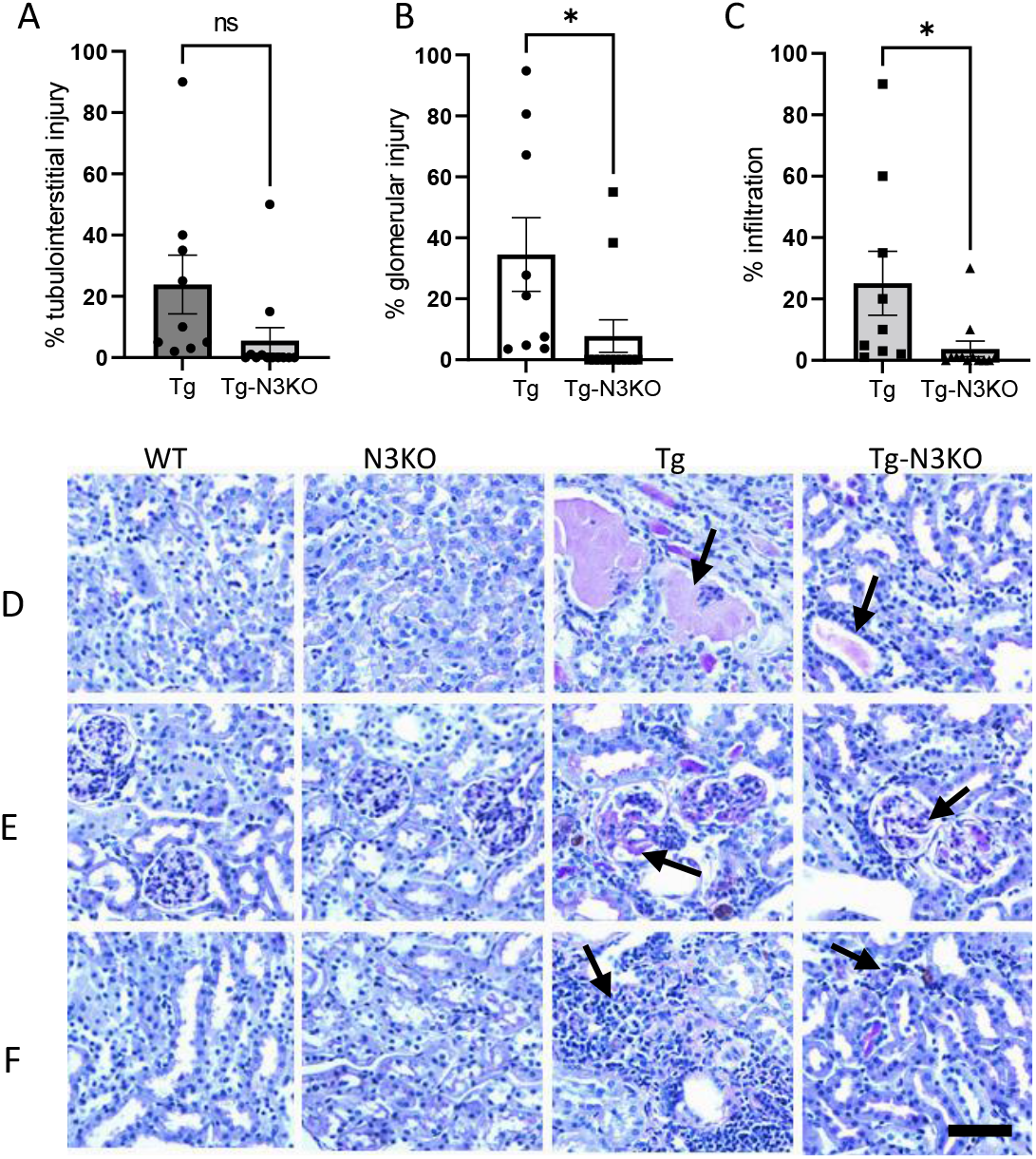
Notch3 deletion ameliorates kidney injury in Tg26 mice. (A, B, C) Kidney sections from 3 months old WT, N3KO, Tg26 and Tg-N3KO mice were stained with Periodic acid Schiff (PAS) to determine kidney injury. PAS staining showing tubulointerstitial injury (a), glomerular tubular injury (b) and percentage inflammation (c). Note severe kidney injury in Tg26 kidneys (n=9) as compared to Tg-N3KO (n=12) kidneys. (D, E, F) Blind quantitation of percent tubulointerstitial injury, percent glomerular injury and percentage infiltration (\**P*<0.05), scale bar 100μm.

### Notch3 deletion inhibits HIV-1

To determine downstream targets of Notch3, RNA sequencing (RNA-seq) was conducted on kidneys of WT, N3KO, HIV-Tg26, and HIV-Tg-N3KO mice. Transcriptome analysis revealed many inflammatory marker genes amongst the topmost upregulated genes in HIV-Tg26 mice (Table:1). Strikingly we also saw that the expressions of HIV-genes nef and env were downregulated in HIV-Tg-N3KO mice compared to HIV-Tg26 mice (Fig: 4A, red asterisks) (Table: 1). These results were confirmed by mRNA expression where a two-fold decrease in both *nef* and *env* was apparent in HIV-Tg-N3KO mice (Fig: 4C, D). Previously, we reported that N4KO in HIV-Tg26 mice also ameliorates disease by reducing inflammation^36^. Thus, to identify differences in expression of *nef* and *env* in N3KO and N4KO in HIV-Tg26 mice, we compared the transcriptomic data from HIV-Tg-N3KO and HIV-Tg-N4KO mice (Table 1). Both HIV-Tg-N3KO and HIV-Tg-N4KO mice had lower *env* and *nef* compared to HIV-Tg26 mice. Quantitative PCR further confirmed these results for *nef* (Fig: 4D). To determine whether the effects on HIV genes are directly related to activation of the HIV-LTR promoter by N3 intracellular domain (N3IC), we conducted promoter reporter assays in podocytes. The N3IC construct was able to increase the LTR promoter activity to 5 folds indicating that *nef* and *env* reduction by N3KO was a direct effect on the HIV-LTR promoter (Fig: 4E). In reverse experiments we infected a human podocyte cell line with either pseudotyped HIV-1 (PNL4) lentiviral vector or its corresponding empty vector (EV). Western blots analysis lysates from these cells demonstrated a significant upregulation of Notch3 intracellular domain (N3IC) (Fig: 4F) revealing a feedback mechanism.

**Fig. 4.**
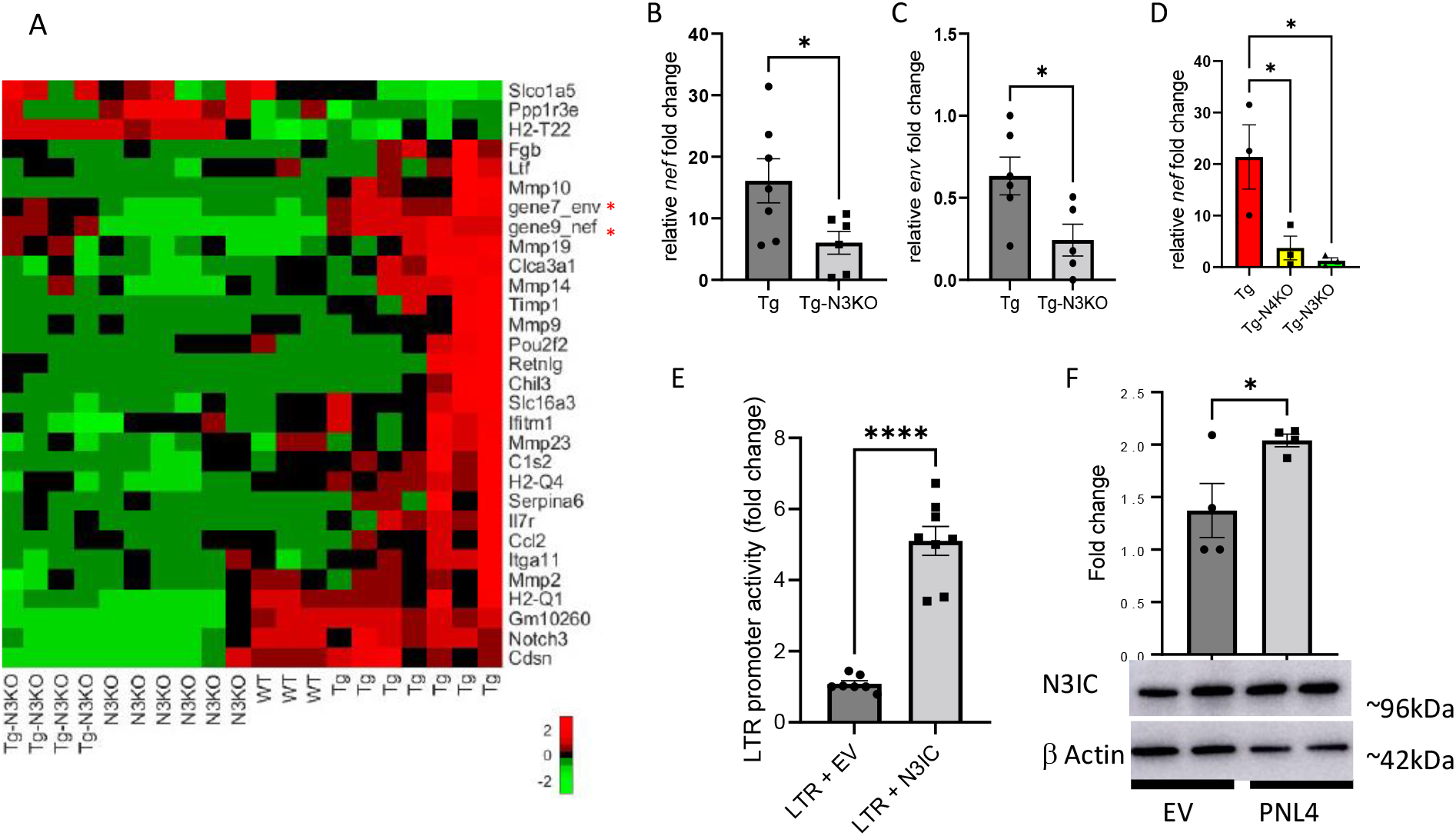
Notch3 targets HIV-1 activity. (A) Heat map showing differential expression of genes obtained from mRNA sequencing of kidneys from 3 months old WT, N3KO, Tg26, Tg-N3KO. Note asterisks for *nef* and *env*. (B and C) Quantitative PCR (qPCR) validating *nef* and *env* (n=5-7) (D) qPCR to compare Tg-N3KO and Tg-N4KO in repressing *nef* (n=3 per condition). (E) HIV-LTR promoter luciferase construct and N3IC or PCDNA3.1 (empty vector (EV)) were transiently transfected in podocytes followed by promoter reporter luciferase assays. Data was averaged from 8 assays in 3 independent experiments. Vector transfected control in each experiment was set to one and relative fold change of LTR promoter activity was calculated. (F) Podocytes were transfected with HIV-LTR construct (PNL4) or EV. After 24 hours, lysates were subjected to western blots for N3IC and β-actin. Data was normalized to β-actin via image J and presented as fold change (n=4).

**Table 1:** Genes downregulated in Tg-N3KO and Tg-N4KO mice compared to Tg26 mice. Genes that were at the top of list for most upregulated genes in HIV-Tg26 verses WT mice were selected and their expression was compared with that of Tg-N3KO or Tg-N4KO. Note that many genes were downregulated significantly as shown by the P values in both Tg-N3KO and Tg-N4KO except *ccl2* and *mmp10* (grey rows) that were only significantly downregulated in Tg-N3KO but not in Tg-N4KO. (FC: fold change)

| *Gene ID* | Tg-N3KO vs Tg26 |  | Tg-N4KO vs Tg26 |  |
| --- | --- | --- | --- | --- |
|  | FC | *P* value | FC | *P* value |
| *chil3* | -133.05 | 0.000931 | -77.2327 | 0.000114 |
| *s100a9* | -17.8874 | 7.51E-05 | -7.92142 | 0.000967 |
| *mmp10* | -14.5953 | 0.001577 | -1.90802 | 0.281182 |
| *Il7r* | -9.05227 | 5.90E-05 | -2.62872 | 0.04056 |
| *ccl2* | -14.9554 | 1.44E-05 | -1.4202 | 0.459143 |
| *retnlg* | -11.5474 | 0.014857 | -12.7715 | 0.004266 |
| *itgam* | -4.48867 | 0.000205 | -2.31992 | 0.012634 |
| *nef* | -2.53002 | 0.00411 | -1.79754 | 0.035797 |
| *env* | -1.64505 | 0.025401 | -1.85543 | 0.001879 |

### Notch3 deletion inhibits invasion of inflammatory cells in HIV-kidneys

Volcano plots derived from RNA seq analysis indicated that differentially expressed genes in HIV-Tg26 verses WT kidneys such as *chil3*, *retnlg*, *ccl2* and *mmp10* skewed towards highly upregulated (Log_2_ (fold enrichment)) in HIV-Tg26 kidneys (Fig: 5A, top). These genes were undifferentiated between Tg-N3KO and WT mouse kidneys (Fig: 5A, bottom). We have previously reported a list of differential gene expressions between HIV-Tg26 and WT mice^37^. Genes that were most significantly downregulated in HIV-Tg-N3KO versus HIV-Tg26 kidneys are shown in Table 1. Raw data was submitted to NIH: sequence read archive (SRA) website with accession numbers, PRJNA578136 (WT and HIV-Tg26), PRJNA680191 (N3KO), PRJNA1010236 (HIV-Tg-N3KO) and PRJNA580295 (N4KO and HIV-Tg-N4KO).

**Fig. 5.**
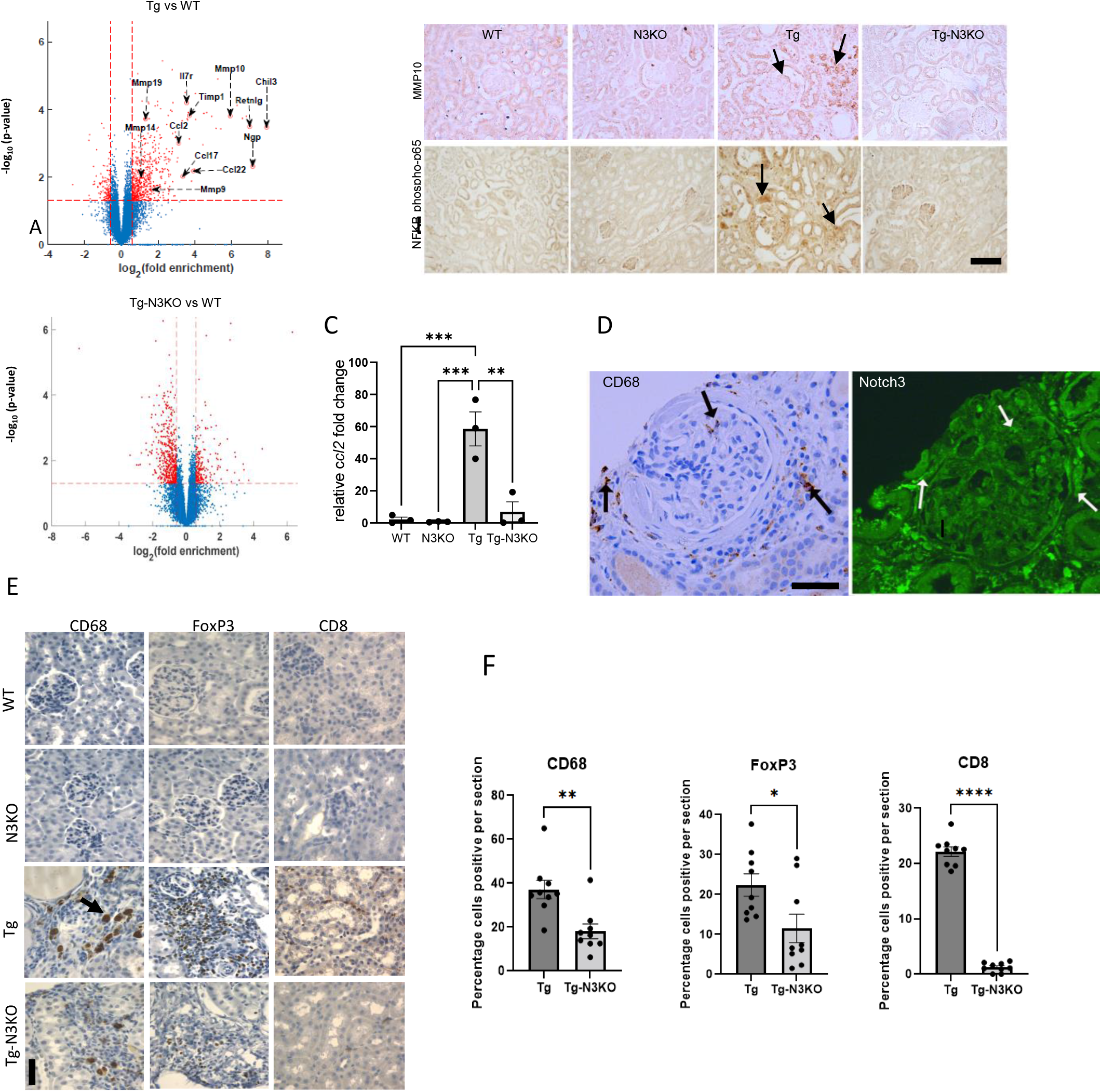
Notch3 targets immune cell infiltration. (A) Volcano plot comparing gene expression in kidneys from Tg vs WT (top) and Tg-N3KO vs WT (bottom) mice. (B) Immunohistochemistry (IHC) for MMP10 and NFkB-p65 in renal sections of WT, N3KO, Tg26 and Tg-N3KO mice. Arrows show glomerular and tubular areas with high expression. (C) quantitative PCR for ccl2 expression. (D) Serial sections from HIVAN patient biopsy immunostained for the presence of CD68 and Notch3. Note the presence of CD68 positive cells in and around glomerulus, the same areas were also labelled bright for the presence of Notch3 (arrows). (Scale bar: 100µm). (E) Paraffin sections from kidneys of WT, N3KO, Tg26, Tg-N3KO mice, labeled for immune cell markers: CD68, FoxP3 and CD8. (F) Quantification of CD68, FoxP3 and CD8 positive cells from Tg26 and Tg-N3KO mice, blindly assessed in Tg26 (n=9) and Tg-N3KO (n=9) kidneys. Each dot represents percent positive cells from the entire kidney where inflammatory invasions were prominent (\**P*<0.05), (Scale bar: 50µm). (\**P*<0.05,\*\**P*<0.01, \*\*\**P*<0.001).

Previously, we reported that N4KO in HIV-Tg26 mice also ameliorates disease by reducing inflammation^36^. Thus, to identify differences between N3KO and N4KO in HIV-Tg26 mice, we compared the transcriptomic data from HIV-Tg-N3KO and HIV-Tg-N4KO mice (Table 1). Compared to HIV-Tg-N4KO mice, HIV-Tg-N3KO mice were more protected against disease as indicated by the robust decrease in the expression of certain inflammatory genes such as *mmp10* (matrix metallopeptidase 10) and *ccl2 (*chemokine (C-C motif) ligand 2)^38–42^. In HIV-Tg26 kidneys, we confirmed the upregulation of MMP10 via immunohistochemistry and *ccl2* via quantitative PCR (qPCR) and also found upregulated NFkB-p65 expression similar to MMP-10 showing inflammatory activity (Fig:5B, C). Further, we validated upregulation of other inflammatory genes known to play a major role in HIV pathogenesis (*tnfa*, *s100a9*, *Itgam*) (Supplementary Fig: S2). Together these data indicate that N3KO alleviates HIV activity and associated inflammation in the kidneys of HIV-1 mice. Next, we asked if N3KO leads to an overall reduction in the number of CD68+ macrophages. CD68+ macrophages were found in clumps in HIV-Tg26 kidneys (Fig: 5E, arrow). Quantification of these cells revealed a 50% reduction in HIV-Tg-N3KO kidneys compared to HIV-Tg26 kidneys (Fig: 5F). In contrast WT and N3KO kidney sections were negative for CD68 (Fig: 5E). To assess other infiltrating cells that N3KO may affect, we conducted staining for FoxP3 (a marker for T Regulatory cells), CD8 (T lymphocytes cells), Granzyme B (found in NK cells and cytotoxic T cells) and CD11c (a marker for dendritic cells). Interestingly, there was a marked increase in the FOXP3 and CD8+ (Fig: 5E) cells, however Granzyme or CD11c+ cells were not found in kidneys (supplementary S3). These data indicate that N3KO reduced the invasion of many inflammatory cell types in the HIV-Tg26 kidneys.

### Notch3 is activated in bone marrow derived macrophages from HIV-Tg26 mice and deletion of *Notch3* alleviates systemic inflammation

Since macrophage markers and infiltrating cells were reduced in HIV-Tg-N3KO kidneys, we reasoned that recruitment of mononuclear cells/macrophages from the bone marrow play a role in the pathogenesis of renal and systemic lesions in HIV-Tg26 mice. To determine whether bone marrow derived macrophages (BMDMs) from HIV-Tg26 mice have active Notch signaling compared to those derived from WT mice, we isolated BMDMs. Cells stained positive for F4/80 marker at differentiation (Fig: 6A and B). Western blots revealed that renal Notch 3IC and Jagged 1 were significantly increased in HIV-Tg26 mice compared to WT controls (Fig: 6C, D, E, F), while delta like 4 (Dll4) did not reach significance. These data reflect that Notch pathway activation in the BMDMs itself may be responsible for inflammation in the kidneys and other organs of these mice. Next, we measured the circulating levels of TNFα, MCP1 to determine if BMDM activation of Notch3 relates to overall inflammation. Our data showed that HIV-Tg-N3KO mice had a significant reduction in the levels of serum TNFα and MCP1 compared to HIV-Tg26 mice (Fig. 6G, H). In addition, dot blot analysis showed reduction in the levels of various cytokines and chemokines in the serum of HIV-Tg-N3KO mice when compared with HIV-Tg26 mice (supplementary S4) (supplementary tables 1 and 2). Collectively, these data suggest dysregulation in the Notch pathway of bone marrow cells in HIV-Tg26 mice, translate into systemic inflammation, and thus, infiltration of immune cells into various tissues. Notch3 deletion ameliorated these defects and the overall phenotype of the HIV-Tg-N3KO mice was improved relative to HIV-Tg26 mice, including a drastic improvement in skin lesions.

**Fig. 6.**
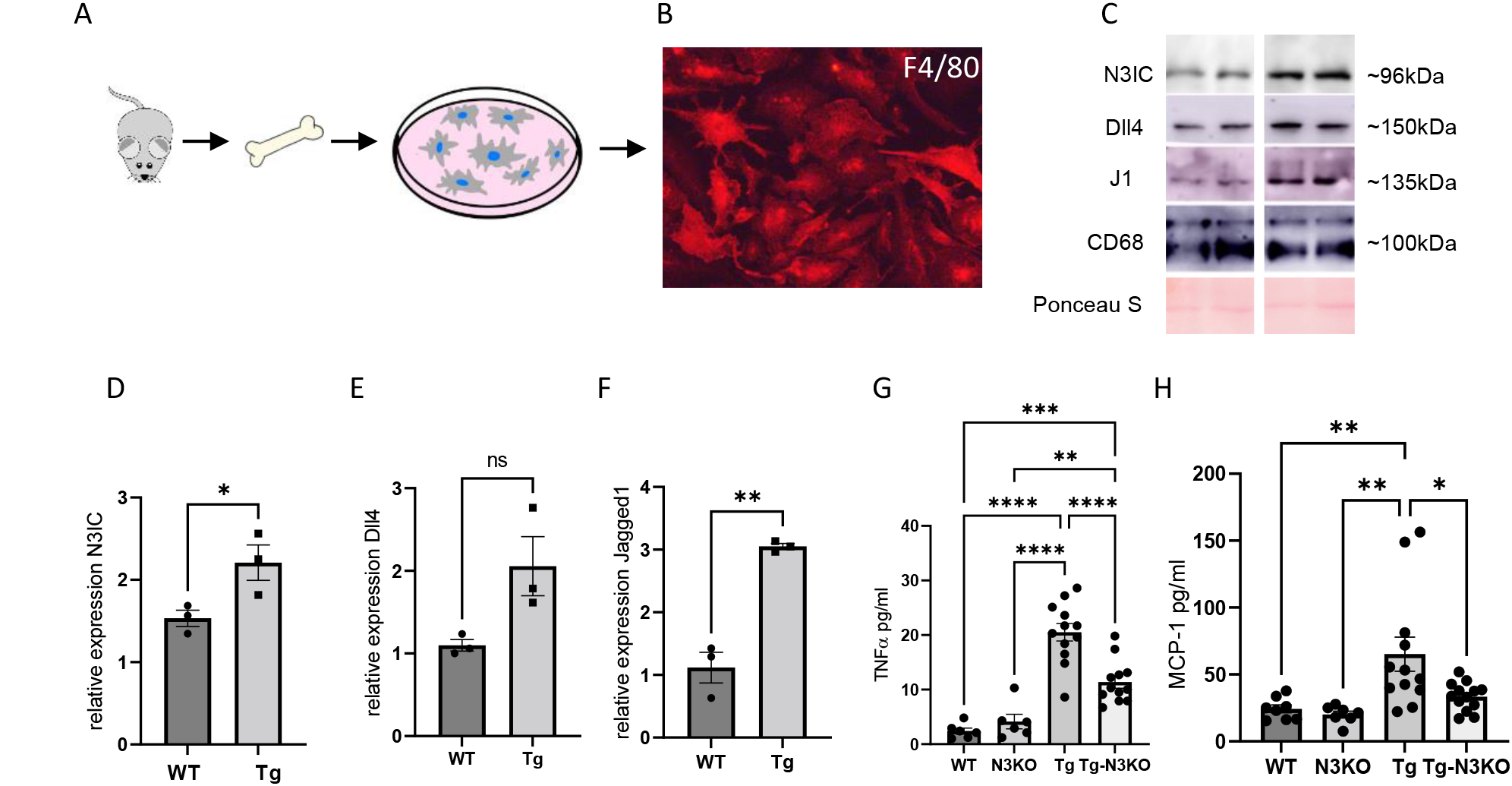
Abnormal Notch3 expression in bone marrow derived macrophages and systemic inflammation is targeted by Notch3 deletion. (A) Schematic showing macrophage isolation from bone marrow in mice. (B) Labeling of differentiated bone marrow cells expressing macrophage marker F4/F80 in fixed cells. (C) Lysates obtained from differentiated macrophages of WT and Tg26 mice were subjected to Western blot analysis for N3IC, Dll4, and Jagged1. CD68 was used as a loading control. (D-F) Quantification of bone marrow cells obtained from 3 mice in each group (n=3). (G and H) ELISA assays were performed for the presence of TNFα and MCP-1 in serum obtained from 3 months old WT, N3KO, Tg26 and Tg-N3KO (n=6-12) mice, both males and females. Data is expressed as pg/ml (\*\**P*<0.01, \*\*\**P*<0.001, \*\*\*\**P*<0.0001).

## Discussion

Our study describes HIV-1 gene regulation and inflammation as a novel dual mechanism by which the Notch3 pathway affects clinical outcomes and survival of HIV-Tg26 mice. Furthermore, to the best of our knowledge, this study demonstrates for the first time that inhibition of Notch3 signaling can specifically improve outcome of HIV-chronic kidney disease, uncovering a potential new target to prevent their progression in people living with HIV-1.

ART is a clinically successful cocktail of antiviral drugs (albeit with side effects) that target various stages of HIV replication cycle. HIV-Tg26 mice carry a 7.4kb proviral HIV-1 DNA construct containing a deletion encompassing most of the *gag* and *pol* genes, and thus HIV-1 cannot replicate in these mice. Regardless of the absence of *gag* and *pol*, the HIV-LTR promoter is fully functional to support the expression of other HIV genes (e.g. *nef*, *env,vpr*) in the HIV-Tg26 mice and cause several lesions, including skin papilloma’s, cataracts and CKD^26,30,43^.

Using the HIV-1 rodent models, we have previously shown that Notch ligands and receptors are activated in a variety of renal cells^7,8^. In previous studies, Notch4 deletion in Tg26 mice resulted in a marked improvement in renal function, an increase in life span and a reduction in renal inflammatory infiltrates. However, the Notch3 expression pattern and its role were not clearly defined^8^. Here, we show that Notch3 was expressed and elevated in both glomerular and extraglomerular regions in HIV-Tg26 mice. A sharp increase of Notch3 expression in cells lining the Bowman’s capsule was observed compared to WT mice. This may signify regeneration potential of PEC progenitors into podocytes and the role of Notch in this phenomenon has been studied in many other kidney injury models^22,44–47^. This area in HIV associated kidney diseases remains a topic of future investigations. N3KO resulted in amelioration of renal disease progression and an increased life span in HIV-Tg26 mice. Compared to N4KO, N3KO resulted in better survival rate of Tg26 mice by 6 months (75% compared to 65%). RNA sequencing data showed macrophage associated inflammatory genes to be upregulated in HIV-Tg26 kidneys, which normalized upon N3KO as compared to N4KO. Matrix-metalloproteinase (MMP) family genes were upregulated in Tg26 kidneys; however, *mmp10* and *ccl2* were uniquely downregulated by N3KO. The MMP proteins either promote or restrict macrophage flux into the tissues^48^. In diabetic kidneys MMP10 contributed to the inflammatory macrophage response which was ameliorated by MMP-10 knockout^49^. In fact, Notch target protein Hey2 has been reported to regulate MMP10^50^. We show that protein expression of MMP10, similar to NFκB (p65), was reduced by N3KO.

The role of Notch3 in inflammation has been studied in non-HIV kidney diseases. Notch3 was activated in lupus nephritis and extra capillary glomerulonephropathy. In nephrotoxic sheep serum (NTS) induced kidney injury and ischemia reperfusion injury N3KO mice exhibited disease protection compared to WT mice. Renal macrophages derived from N3KO mice failed to activate inflammatory cytokines^23^. *In vitro*, overexpression of active Notch3 in podocytes led to reorganized cytoskeleton leading to a migratory, proliferative and proinflammatory phenotype^22^. Since there was less inflammatory infiltration in HIV-Tg-N3KO mice compared to Tg26 mice, we reasoned that it may be due to less infiltrating cells from extrarenal sources. The RNA seq supported our reasoning showing a drastic decrease in the macrophage associated markers in the HIV-Tg-N3KO kidneys as opposed to HIV-Tg26 kidneys. Thus, we focused on the BMDMs. To our surprise, we found activation of Notch3 signaling and upregulation of Notch ligands in the BMDMs from HIV-Tg26 mice. Similar to previous studies,^23^ we speculate that Notch3 activation in combination with HIV genes in glomerular cells and BMDMs may activate the migratory, proliferative and infiltrating properties of cells. We also speculate that intraperitoneal macrophages or cells of other lineage such as Tregs (FoxP3 positive) may also have Notch3 activation in the HIV-Tg26 mice. Taken together, these changes may be responsible for the renal and systemic inflammatory responses and thus N3KO led to decreased systemic TNF alpha and MCP1, as well reduced recruitment of inflammatory cells in the kidney and likely in skin lesions, thereby drastically increasing the life span of these mice.

Since the Notch pathway is activated in glomerular diseases with or without HIV-1, it remains to be determined whether HIV-1 can induce the activation of Notch3 directly. Using a pseudotyped replication defective HIV-1 construct, we found that HIV-1 can activate Notch3 directly in podocytes. In addition, N3KO in HIV-Tg26 mice resulted in a significant drop in the expression of *nef* and *env* and N3IC significantly activated the HIV-LTR promoter activity indicating a feedback regulation of both N3IC and HIV-1 genes. In support of these findings, the HIV-Tat protein was shown to interact with the extracellular domain of the Notch receptors and was suggested as a non-canonical ligand for Notch activation^51–54^. In addition, previous studies in HIV-Tg26 mice and human podocytes showed that HIV-1 can precipitate the development of HIVAN and affect the cytoskeletal structure of podocytes cultured from children with HIVAN^55,56^. Moreover, there are two RBP-JK binding sites adjacent to the NFκB binding sites on the HIV-LTR promoter. RBP-JK binds the LTR and promotes recruitment of histone deacetylate (HDACs) and polycomb repressive complexes (PRC) in CD4+ T cells. Knock down of RBP-JK or PRC was shown to result in proviral activation^57–59^. RBP-JK is a known transcriptional repressor which changes into a transcriptional activator upon NIC binding. Thus, we speculate that NIC binding to RBP-JK in HIV-Tg26 leads to activation of pathways downstream of Notch3 in HIV-Tg26 mice. Further studies will be required to clarify this mechanism, since, in HIV-Tg26 mice, the LTR promoter is primarily bound by the active p50/p65 heterodimer ^60^.

While Notch3 inhibition appears to be an excellent strategy to reduce inflammation and HIV gene expression, we stress that a combined approach to decrease the activity of both Notch 3 and Notch4 may have better outcomes. However, since age related vascular lesions have been detected in Notch3 mice ^61,62^, targeted delivery of Notch3 inhibitor may be beneficial.

We have found that the Notch3 pathway is activated in BMDMs and several kidney cell types from HIV-Tg26 mice. Activation of the Notch3 pathway leads to significant systemic and renal inflammatory changes that precipitate the development of chronic kidney disease and affect the survival of these mice. Our data also suggest that HIV-1 genes can directly activate the Notch3 pathway in immune cells and human podocytes. It is also tempting to speculate that the Notch3 pathway may interact with other cells of the innate immune system as well as major risk factors that predispose to the development of CKDs in people of African descent, including the APOL1 risk variants. Nonetheless, regardless of the APOL1 risk variants, it appears that new therapies that target the Notch3 pathway, may have a key role preventing the development of HIV-CKDs by decreasing HIV-1 genes and inflammation in kidneys and likely other organs.

## Material and Methods

### Sex as a biological variable

Human biopsy samples are unidentified, age matched and consist of both males and females. In mouse studies both males and female were included in equal numbers and no difference in sexes was apparent.

### Human tissues

Renal biopsies were from patients with HIVAN, and age matched normal controls (both males and females) were used. The use of these tissues for the study was evaluated and approved for use by the Human Protection Program at the University of Kansas Medical Center and University of Feinstein Institute for Medical Research, Zucker School of Medicine at Hofstra-Northwell.

### Animal care and studies

All experiments were performed under guidelines according to the Guide for the Care and Use of Laboratory Animals of the National Institutes of Health, and as approved by the Institutional Animal Care and Use Committee of the University of Kansas Medical Center (Kansas City, KS, USA). Mice were housed in micro-isolator cages on a high-efficiency particulate air-filtered, ventilated rack under aseptic and pathogen free conditions. Animal breeding strategies, renal function evaluation and histological protocols can be found in supplementary methods. The heterozygous HIV-Tg26 mice show all characteristics of HIVAN and were a kind gift from Dr. Paul Klotman (Baylor College of Medicine, Houston, TX)^28^. Notch3^d^^1^ [N3KO] mice were obtained from the Jackson Laboratory (JAX:023807). All mice were maintained on FVB/N background. Sample size was based on power analysis and previous studies^36^. All experiments are performed/reported under ARRIVE guidelines^63^ and according to the Guide for the Care and Use of Laboratory Animals of the National Institutes of Health, and as approved by the Institutional Animal Care and Use Committee of the University of Kansas Medical Center. Mice were housed in micro-isolator cages on a high-efficiency particulate air-filtered, ventilated rack. Conditions were aseptic and pathogen free. Comparisons were made between Wildtype (WT, normal control), HIV-Tg26 (positive control), N3KO mice (another control) and HIV-Tg-N3KO mice (new genotyping to be tested) at 3 months of age. N3KO mice (B6;129S1) were backcrossed onto FVB/N background for at least six generations to be congenic before breeding with HIV-Tg26 mice. Before euthanasia at 3 months of age, overnight urine was collected in metabolic cages for proteinuria and albumin-to-creatinine ratio evaluation. Blood was collected at the time of euthanasia for serum isolation. One half of a kidney was stored in paraformaldehyde overnight and changed to 70% ethanol. The other half and second kidney were snap frozen for RNA isolation and protein lysates and stored at −80C immediately. Another set of mice were monitored for 6 months for mortality studies.

### Antibodies and reagents

The following antibodies were used: anti-β-actin (1:1000; Sigma-Aldrich, St. Louis, MO, USA; A5441); anti-MMP-10 (1:1000; My Bio Source, San Diego, CA, USA; MBS2027749); anti-Notch3, DLL4 and Jagged1 (1:1000; Abcam, Cambridge, UK; ab23426, ab7280 and ab7771, respectively). Anti-CD68 for detection in mice (1:100, Abcam, Ab125212); anti-CD68 for detection in human (1:100; BioCare Medical, CM033A); anti-p-NF-κB (Cell Signaling Technologies, 3033s), Mouse immune Cell phenotyping IHC antibody kit (from Cell Signaling Technologies (37495) containing anti-FoxP3, CD11c, CD8 and Granzyme B antibodies); secondary antibodies goat anti-mouse IgG H&L (Alexa Fluor® 594) (ab150116) and goat anti-rabbit IgG H&L (Alexa Fluor® 488) (ab150077), both from Abcam.

### Renal function evaluation

Blood urea nitrogen levels were quantified using Quantichrom Urea Assay Kit (Bioassay Systems, Hayward, CA) according to manufacturer’s protocol. Proteinuria was measured in 2 µL of urine which was loaded onto 10% SDS-polyacrylamide gel for electrophoresis, followed by Coomassie blue staining. Bovine serum albumin (BSA) (2 µl; 10 mg/ml) was used as a positive control. Albumin in urine was assessed by ELISA kit from Bethyl Labratories (Houston, TX). Creatinine levels in same urine samples were quantified using QuantiChrome Creatinine Assay kit (Bioassay Systems).

### Histology

Kidneys were fixed in 4% paraformaldehyde overnight and transferred to 70% ethanol. The tissues were processed and embedded in paraffin at the core facilities at the University of Kansas Medical Center. Five-micrometer sections were stained with Hematoxylin & Eosin, periodic-acid Schiff (PAS) and Mason Trichrome (MTA). Slides were examined blindly and scored for tubulointerstitial disease and glomerular injury (n=6, each group). The area was inspected and percentage of tubulointerstitial disease (enlarged, reactive tubular nuclei, tubular casts, tubular dilation and interstitial fibrosis) was estimated. The number of glomeruli with visible disease (segmental and global sclerosis, collapsed phenotype, adhesion to Bowman’s capsule) was counted for glomerular injury score. Percent immune cell infiltration was calculated by estimating the percent area affected cortex.

### Cell culture

All cell lines used were free of mycoplasma contamination, as verified by Lonza kit (LT07-118). Human immortal podocytes were kind gift from Dr. Moin Saleem (University of Bristol, UK) ^64^. Immortal podocytes were maintained in a growth media containing RPMI 1640 (Hyclone) [supplemented with 10% FBS, 1× Pen/Strep, 1× insulin-transferrin-selenium (GenDEPOT)] at 33°C for SV40T antigen expression. To inactivate SV40T antigen, cells at 50% confluency were moved to 37°C in 5% CO2 and allowed to differentiate for 7-10 days. Description of podocyte infections with lentiviral constructs for Notch3 induction and bone marrow derived macrophage isolation and culture are described in supplemental material and as described^65,66^.

#### Podocyte transductions

To study the induction of Notch3 by HIV-1, pNL4-3:ΔG/P-GFP construct was used. Generation of this construct has been described before^65,66^. Briefly, in HIV-1 proviral construct (pNL4-3) the *gag*/*pol* regions were substituted with green fluorescence protein (GFP) reporter gene. As a negative control (WT), pHR-CMV-IRES2-GFP-ΔB construct containing HIV-1 LTR and GFP empty expression vector was used. These constructs were pseudotyped with VSV.G envelope separately to generate viral particles in 293T cells. Viral particles were used to infect differentiated conditionally immortalized podocytes for 2 days as described^36^. Podocytes were lysed using RIPA buffer [137 mM NaCl, 50 mM Tris HCl pH 7.5, 12 mM EDTA, 1% IGEPAL and complete protease inhibitor (Thermo Fisher Scientific, Saint Louis, MO)] and stored in −80^0^C until used.

#### Luciferase promoter reporter assays

The HIV-LTR promoter reporter luciferase construct is described previously and obtained as a kind gift from Dr. Wang^67^. N3IC expressing construct was developed as previously reported^68^. Co-transfections were carried out with empty vector (PCDNA3.1), N3IC construct (PCDNA3.1 N3IC) and HIV-LTR (pGL3-LTR-Luc) using [Lipofectamine™ LTX](https://www.thermofisher.com/order/catalog/product/A12621) (Thermo Fisher). 24hrs after transfections, cells were lysed with passive lysis buffer and Dual luciferase assay kit (Promega) was used to assess luciferase activity using luminometer. Reporter activity was normalized to renilla luciferase activity and expressed as fold change of relative light units.

#### Isolation and culture of bone marrow derived Macrophages (BMDMs)

Mice were euthanized, and legs were removed under aseptic conditions. Legs were deskinned and tibia and femurs were separated. Both ends of the tibia and femur were cut under sterile conditions and marrow was flushed by placing femur or tibia on a cut pipette tip placed into a 1.5 mL tube. The tubes were then spun down at 250g for 5 minutes at room temperature to pellet cells. The cell pellet was resuspended in tissue culture plate with BMDM media containing IMDM media supplemented with glutamine, 1% minimum essential medium (MEM) non-essential amino acids, 1% sodium pyruvate, 1% penicillin/streptomycin, 0.05mM b-mercaptoethanol and 30% L929 supernatant for a source of macrophage colony stimulating factor (see below). Cells were incubated for 7 days at 37^0^C in 5% CO_2_ with addition of 5ml/10cm2 dish media at day three. On day eight, the cells were scraped gently and in PBS, resuspended and counted. 2X10^6^ cells were plated in 10cm^2^ dishes for three days and lysed. Lysates were stored in −80^0^C until used.

#### L929 media

L929 cells were grown to confluency in Iscove’s Modified Dulbecco’s *Medium* media containing 10% fetal bovine serum, 1% non-essential amino acids, sodium pyruvate and 1× Pen/Strep. After 3 days media was collected, filtered sterilized and stored at −80°C until use.

### Immunolabeling

Sections from kidneys of WT, N3KO, Tg, and Tg-N3KO mice were deparaffinized with xylene and hydrated in ethanol. These sections were then boiled in sodium citrate buffer (10 mM sodium citrate, 0.05% Tween 20, pH 6.0) and cooled to room temperature. To block endogenous peroxidase activity, sections were incubated for 30 min with 3% hydrogen peroxide. They were then washed in PBS and blocked with 10% normal serum (from species the secondary antibody was raised in, in PBS) for 1 h, followed by a 1 h incubation in a humidified chamber with primary antibodies. Sections were washed three times with PBS, then incubated for 1 h in 1:400 diluted biotin-conjugated secondary antibodies (Vector Laboratories) for IHC and fluorescein/Texas Red-conjugated antibodies for immunofluorescence (IF), followed by another wash. For IF, the slides were mounted using Alexa Fluor® 594 or Alexa Fluor® 488 with DAPI (Abcam). For IHC analysis, the slides were further incubated with avidin-biotin-peroxidase complex (ABC Elite, Vector Laboratories), which was detected with diaminobenzidine (DAB, Sigma-Aldrich). Some kidneys were counterstained with hematoxylin. Tissue sections were then dehydrated with graded ethanols and mounted with permount (Fisher Scientific, Pittsburg, PA, USA). Slides were viewed on a Leica DMLB 100s upright microscope.

### Quantification of histology and immunolabelling

For counting CD68, FoxP3, CD8 labelled cells, images were captured from all the areas where these cells were present (n=5-6 per kidney), 6 kidneys per group). Cells positive for Hematoxylin (total cells in the section) and cells with DAB staining (only positive for CD68, CD8 or FoxP3) were counted and percentage of positively labeled cells was calculated per section and averaged. For quantification of immunofluorescence intensity, image J was used, and percentage intensity was measured from each image and averaged. For quantification of Notch3 intensity, vascular Notch3 values were subtracted from total intensity to assess glomerular and tubular labelling individually.

### Western blots

Protein estimation was performed in cell lysates by BCA protein assay kit (Bio-Rad, Hercules, CA, USA). For Western blots, 75 μg protein was solubilized in 4× NuPage (Novex) sample buffer [containing 25% tris (2-carboxyethyl) phosphine (TCEP)]. Samples were heated to 65°C for 10 minutes before electrophoresis on 10% SDS-polyacrylamide gels. Samples were then transferred to polyvinylidene difluoride (PVDF) membranes. 5% non-fat dry milk in Tris-Buffered Saline containing 0.1% Tween 20 (TBST) was used for blocking of immunoblots for 1 h at room temperature. After washing blots with TBST, overnight incubation was performed using the appropriate dilutions of primary antibodies. Blots were washed three times with TBST, then incubated for 1 h at room temperature in secondary antibodies (Vector Laboratories) (1:5000 dilution in blocking solution). Chemiluminescence was then used for detection (Western Lightning Plus ECL, Perkin Elmer).

### RNA-sequencing

RNA-sequencing was carried out in kidneys from 3 months old WT (n=3), HIV-Tg26 (n=7), N3KO (n=4) and HIV-Tg-N3KO (n=6). Briefly, mouse kidneys were lysed in Trizol (Thermo Fisher Scientific, Saint Louis, MO) using manufacturer’s protocol. Verification of RNA integrity, global transcriptomic analysis, preparation of RNA-seq libraries, sequencing of cDNA libraries was performed using Affymetrix Clariom D (Thermo Fisher Scientific, Saint Louis, MO) array at the Genomics Core Facility of the University of Kansas Medical Center as described before^37,69^. Illumina NovaSeq 6000 sequencing machine (Illumina, San Diego, CA) was used to perform RNA sequencing at a strand specific 100 cycle paired-end resolution. The read quality was assessed using the FastQC software^70^. On average, the per sequence quality score measured in the Phred quality scale was above 30 for all the samples. The sequenced reads were mapped to the combined mouse (GRCm38.rel97) and Human immunodeficiency virus 1 (GCF_000864765.1) genomes using STAR software (version 2.6.1c)^71^. On average, 95% of the sequenced reads mapped to the genome, resulting between 27 and 32 million mapped reads per sample, of which on average 86% were uniquely mapped reads. Transcript abundance estimates were calculated using the featureCounts software^72^. Expression normalization and differential gene expression calculations were performed in DESeq2 software^73^ to identify statistically significant differentially expressed genes. The significance p-vales were adjusted for multiple hypotheses testing by the Benjamini and Hochberg method^74^ establishing a false discovery rate (FDR) for each gene. The resulting data were used to generate heat maps and volcano plots for visualization.

### Quantitative RT-PCR

Total RNA was isolated from tissues using Trizol (Thermo Fisher), following the manufacturer’s protocol. Nanodrop analysis was used to determine RNA concentration and purity. Using a High-Capacity Reverse Transcription Kit (Thermo Fisher Scientific), complementary DNA (cDNA) was prepared from 2 µg total RNA. Quantitative RT-PCR was performed using Power SYBR Green Master Mix (Applied Biosystems). Results were normalized using 18S ribosomal RNA (rRNA) expression and calculated using comparative ΔΔCT. Ct values for each gene were obtained. By subtracting the housekeeping gene value from the Ct value of the samples, delta Ct (ΔCT) values were then calculated. Using the ΔCt values of the WT controls (n=3), an average was found. The average ΔCt value of the WT controls was then subtracted from the ΔCt values of all samples to obtain a ΔΔCt value. To calculate relative fold change, the 2-ΔΔCt formula was used for each control or gene of interest value. Values from each group were averaged and shown as a relative fold gene expression change as previously shown. The primers used in this study are listed in supplementary table 3.

### Enzyme linked immunoassay (ELISA) and chemokine/cytokine profiling

Serum was assessed for presence of TNFα and MCP-1 using mouse ELISA kits (R&D SYSTEMS, USA) according to manufacturer’s instructions. For analysis of chemokines and cytokines in serum of Tg26 and Tg-N3KO mice, serum samples from 3 male mice (200ul each) were pooled from each group. From each group 150 ml serum was used for chemokine array using the chemokine Proteome Profiler Array kit (Cat No# ARY020, R&D SYSTEMS) and 200ul was used from each group for cytokine array using the cytokine proteome profiler array kit (Cat No# ARY028, R&D SYSTEMS). Manufacturer’s instructions were followed.

### Statistics

Data re plotted as mean± standard error (SE). Unpaired Student’s *t*-test was used to measure statistical significance between control and test groups. To compare more than two groups, a one-way ANOVA followed by Tukey’s multiple comparison test was performed, using Graphpad. *P*<0.05 was considered statistically significant.

### Data Availability

The RNA sequencing data has been submitted to Sequence read archives (SRA) and can be availed by using accession numbers, PRJNA578136 (WT and HIV-Tg26), PRJNA680191 (N3KO), PRJNA1010236 (HIV-Tg-N3KO) and PRJNA580295 (N4KO and HIV-Tg-N4KO).

## Author contributions

M.T., N.S., M.M., J.D., S.Y., A.C1., A.R., H.E., T.F and M.S. performed experiments. S.G. A.C2. and P.C performed RNA seq analysis and data sharing. J.T., provided expertise in BMDM isolation, P.R., and N.D performed inflammatory marker assessment. P.S., and P.T., provided resources. M.T., provided expertise in promoter reporter assays. H.E., N.S., and M.S., wrote the manuscript. P.T., M.T. T.F., P.S., P.R., N.D and M.S. critically revised the manuscript.

## Acknowledgements

**Funding**: This publication was supported by National Institute of digestive and diabetic and kidney diseases R01DK108433, an Institutional Development Award (IDeA) from the National Institute of General Medical Sciences of the National Institutes of Health (NIH) under grant number P20 GM103418 and an institutional Lied grant to MS and NIH grant R01 HL1528322 to ND. We thank Clark Bloomer and the genomics core staff at the University of Kansas Medical Center for RNA sequencing which is supported by Kansas Intellectual and Developmental Disabilities Research Center (NIH U54 HD090216), the Molecular Regulation of Cell Development and Differentiation-COBRE(P30 GM122731-03)-the NIH S10 High-End Instrumentation Grant (NIH S10OD021743) and the Frontiers CTSA grant (UL1TR002366).

