## supplemental figures for "Notch3 deletion regulates HIV-1 gene expression and systemic inflammation to ameliorate chronic kidney disease"

Supplementary Figure S1

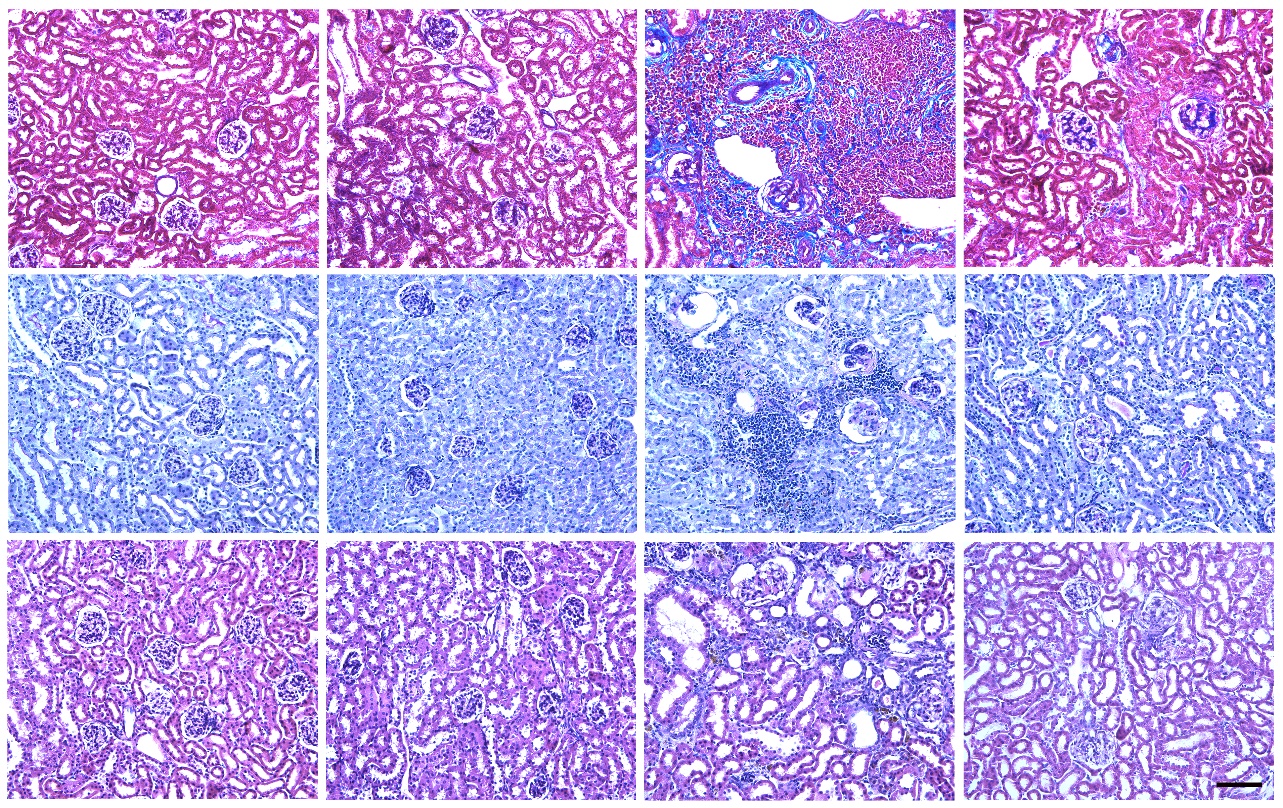

WT	N3KO	Tg26	Tg-N3KO

A

B

C

**Notch3 deletion ameliorates histological lesions in Tg26 mice:** (A,B,C) Kidney sections from 3 months old female WT, N3KO, HIV-Tg26 and HIV-Tg-N3KO mice were stained with Mason Trichrome (MTA), Periodic acid Schiff (PAS) and Hematoxylin & Eosin (H&E) respectively. Representative image shown from each group. Note massive glomerulosclerosis, fibrosis, tubulointerstitial injury and infiltration in Tg26 sections which was reduced in HIV-Tg-N3KO sections. (D) quantification of MTA represented as percent area stained for MTA. (scale bar 50μm), kidneys ( ****P*<0.0001).

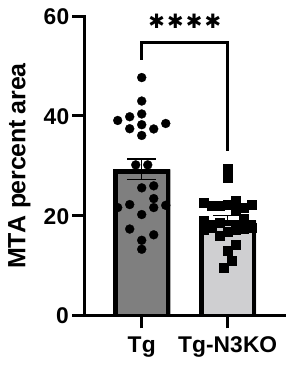

D

Supplementary Figure S2

A

B

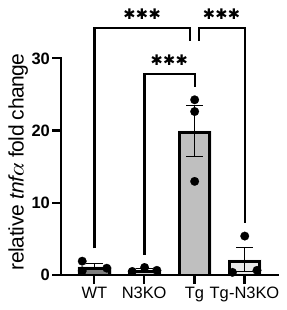

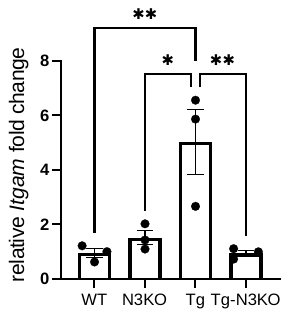

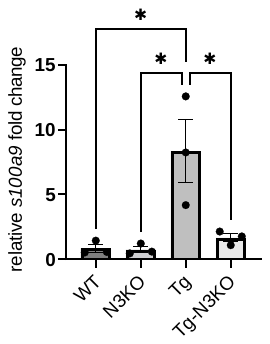

C

**Quantitation of major genes downregulated in Notch3 deleted Tg26 mice.** (A, B, and C) Quantitative PCR validating the upregulated expression of *tnfa*, *itgam* and *s100a9*, genes related to macrophage associated inflammation in Tg26 mice that almost normalized in Tg-N3KO kidneys (**P*<0.05, ** *P*<0.01, *** *P*<0.001).

Supplementary Figure S3

Granzyme

CD11C

WT	N3KO	Tg26	Tg-N3KO	control (mouse spleen)

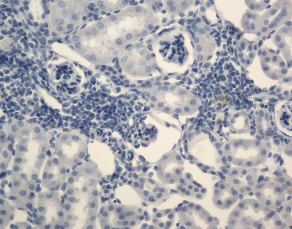

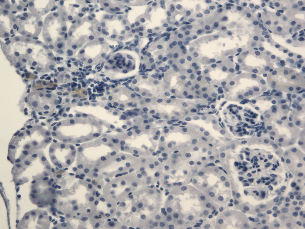

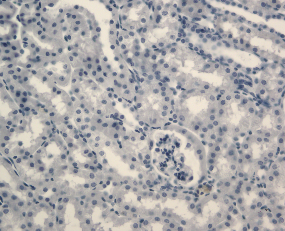

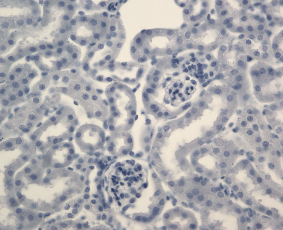

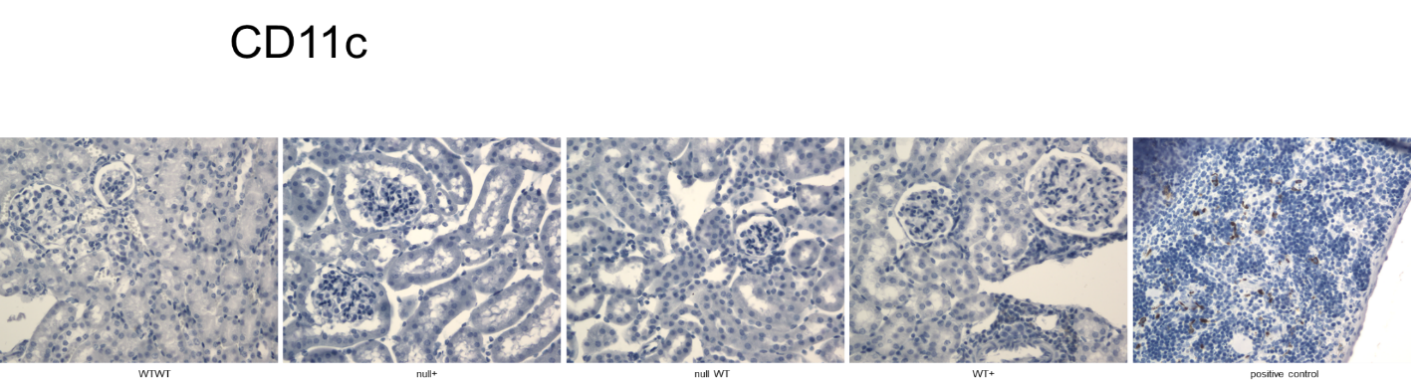

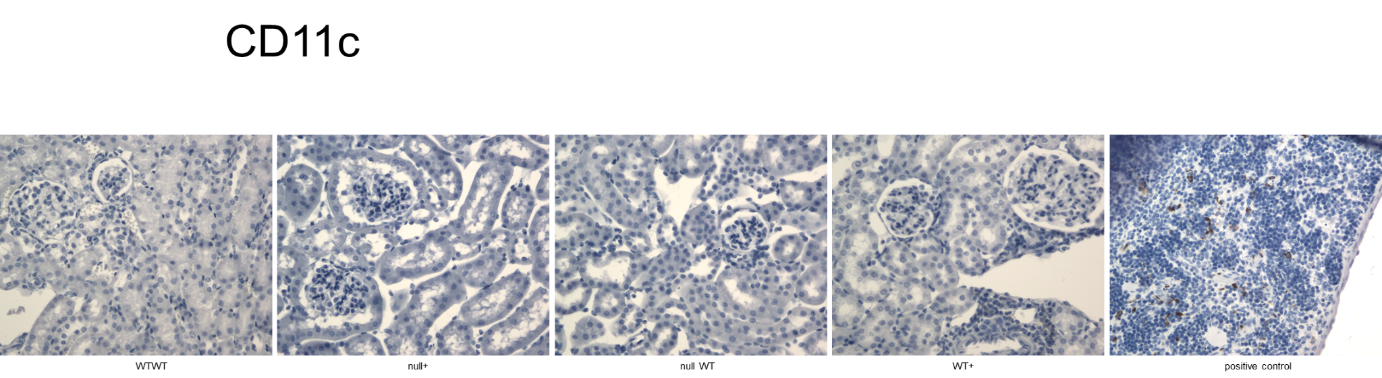

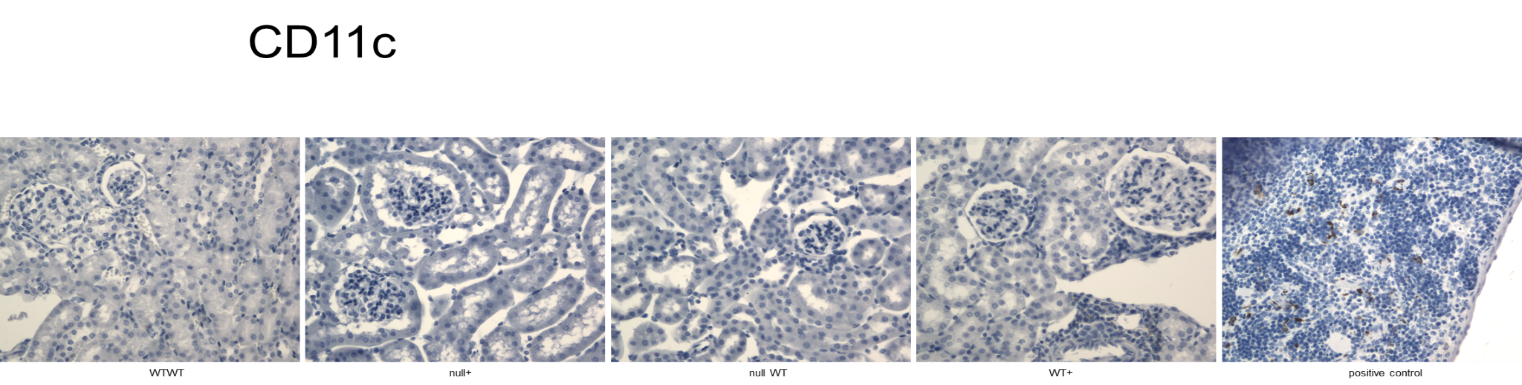

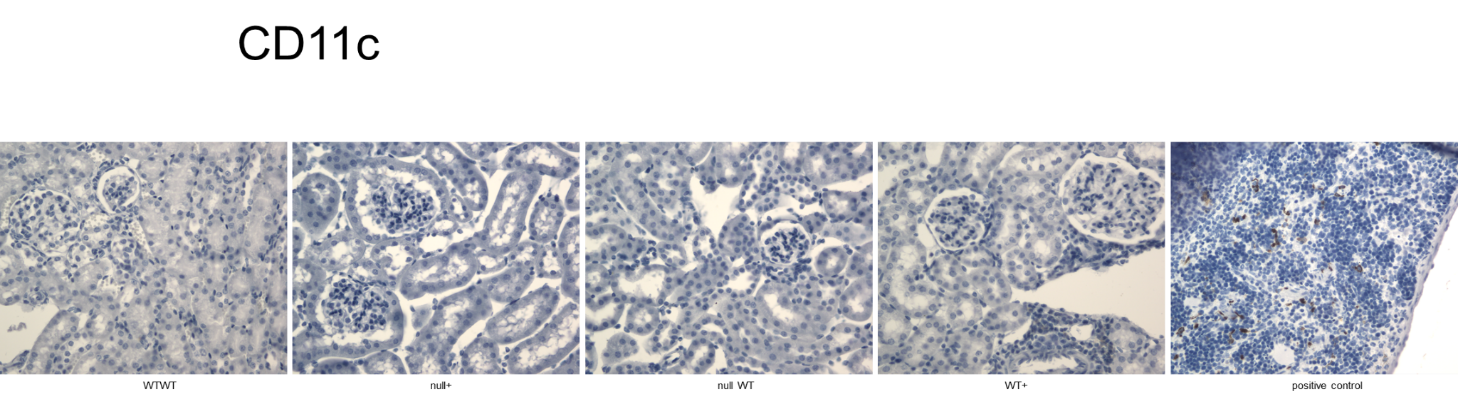

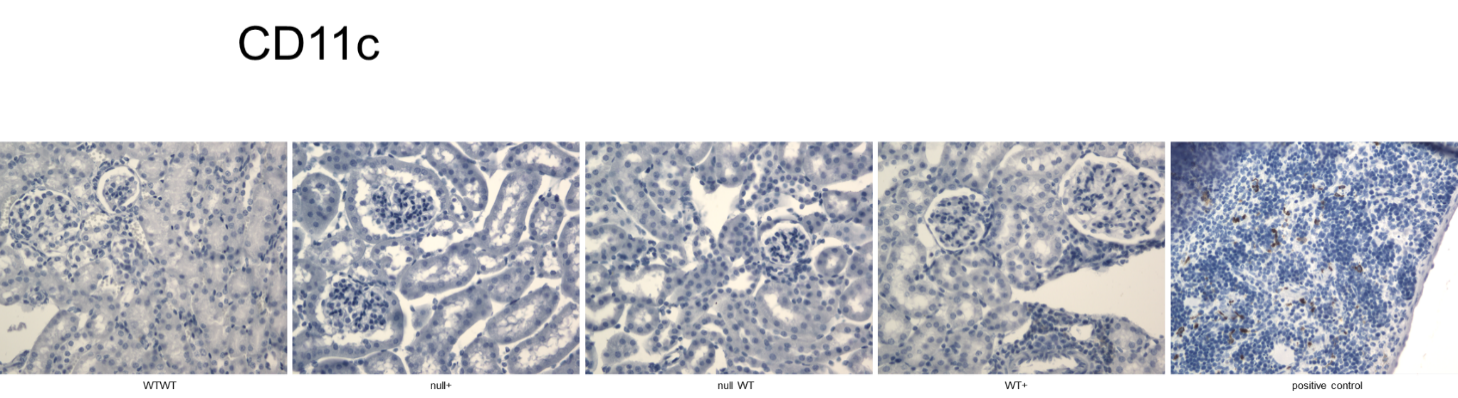

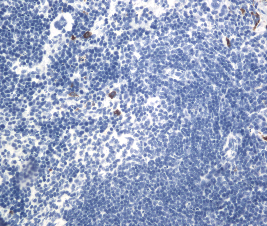

**Effects of Notch3 deletion on Granzyme and CD11C in kidneys of Tg26 mice**. Immunohistochemistry was performed in paraffin sections from WT, N3KO, Tg26 and Tg-N3KO kidneys for presence of Granzyme and CD11C. Data is representative of 4-5 kidneys from each group of male and female mice. Control represents spleen of a normal mouse where positive labeling is noticed (arrows). Scale bar: 100μm.

Supplementary Figure S4

A

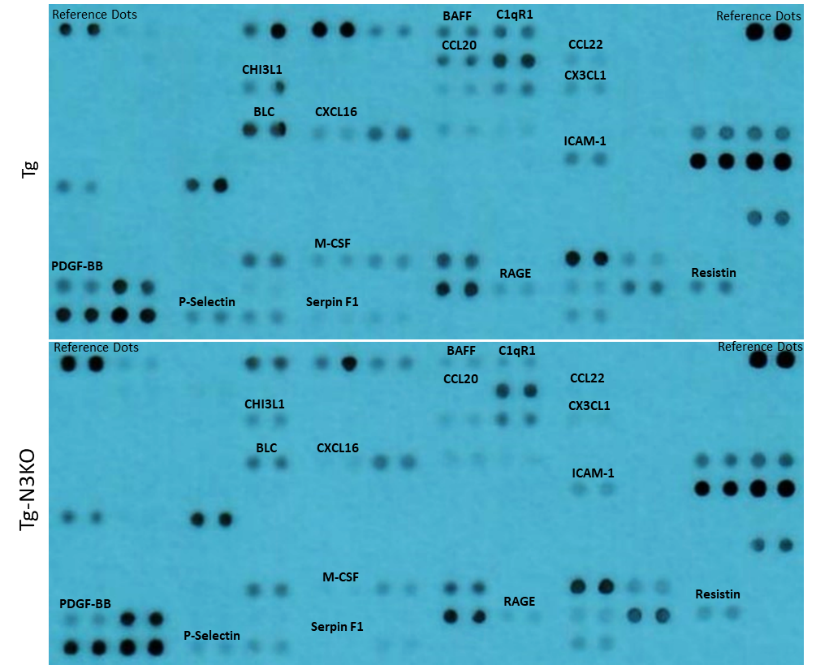

Tg-N3KO

Tg-N3KO

WT

WT

B

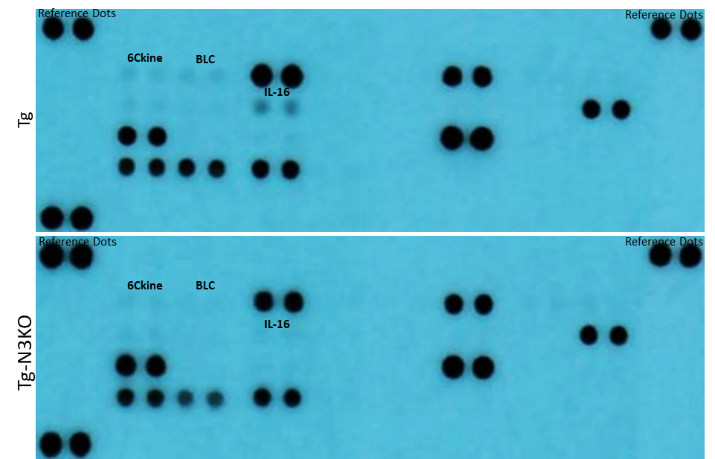

**Cytokine and chemokine regulation by Notch3.** (A and B) Dot blots comparing the expression of various cytokines (A) and chemokines (B) in the pooled serum from Tg26 (n=3) and Tg-N3KO (n=3) mice (males). Dots in duplicates represent one analyte. Dots having ≥ 1.5-fold differences in mean pixel density between Tg26 and N3-TgKO are labelled.
