## supplemental tables for "Notch3 deletion regulates HIV-1 gene expression and systemic inflammation to ameliorate chronic kidney disease"

| **Supplementary Table 1**  **Supplementary Table1**  **List of differentially expressed cytokines.** |  |
| --- | --- |
| Name of Analytes | Tg vs Tg-N3KO |
| Angiopoietin-1 | -1.14 |
| Angiopoietin-2 | -1.34 |
| BAFF/BLyS/TNFSF13B | -1.61 |
| C1q R1/CD93 | -1.92 |
| CCL20/MIP-3 alpha | -6.61 |
| CCL22/MDC | -3.53 |
| CD14 | -1.38 |
| Chitinase 3-like 1 | -1.65 |
| CX3CL1/Fractalkine | -1.58 |
| CXCL13/BLC/BCA-1 | -1.83 |
| CXCL16 | -3.27 |
| ICAM-1/CD54 | -1.46 |
| M-CSF | -8.75 |
| PDGF-BB | -1.48 |
| Periostin/OSF-2 | -1.45 |
| RAGE | -1.90 |
| Resistin | -1.71 |
| P-Selectin/CD62P | -1.97 |
| Serpin E1/PAI-1 | -1.26 |
| Thrombopoietin | -1.58 |
| DPPIV/CD26 | -1.72 |
| Endoglin/CD105 | -4.60 |
| VEGF | -1.37 |
| WISP-1/CCN4 | -2.20 |
| Serpin F1/PEDF | -2.28 |
| CCL6/C10 | +1.91 |
| Myeloperoxidase | +1.04 |
| Osteopontin (OPN) | +1.25 |
| Osteoprotegerin/ TNFRSF11B | +1.10 |
| Pentraxin 2/SAP | +1.30 |
| RBP4 | +1.12 |
| Reg3G | +1.62 |

Supplementary table 1 showing the list of cytokines altered in the serum of Tg-N3KO mice compared to Tg26 mice (samples pooled from n=3, each group).

**Supplementary Table 2**

**List of differentially expressed chemokines.**

| **Name of Analyte** | **Tg vs Tg-N3KO** |
| --- | --- |
| 6Ckine/CCL21/SLC/Exodus-2 | -1.69 |
| BLC/CXCL13/BCA-1 | -2.08 |
| C10/CCL6/MRP-1 | -1.36 |
| Chemerin/RARRES2 | -1.10 |
| CTACK/CCL27/ALP/ILC/Eskine | -1.23 |
| Fractalkine/CX3CL1 | -2.04 |
| IL-16 | -3.65 |
| JE/CCL2/MCP1 | -1.30 |
| LIX/GCP-2/ENA-78 | -1.06 |
| MCP-5/CCL12 | -1.24 |
| MDC/CCL22/ABCD-1 | -1.12 |
| MIG/CXCL9/CRG-10/CMK | -1.07 |
| RANTES/CCL5/SISd | -1.33 |
| CXCL16/SRPSOX | +1.43 |
| Eotaxin/CCL11 | +1.14 |

MCP2/CCl8/HC14	+1.21

Supplementary table 2 showing the list of chemokines altered in the serum of Tg-N3KO mice compared to Tg26 mice (samples pooled from 3 mice in each group).

**Supplementary Table 3**

| Gene | Forward primer (5'-3') | Reverse primer (5'-3') |
| --- | --- | --- |
| *mmp10* | GCCCAGCTAACTTCCACCTTT | GAGAGTGTGGATCCCCTTTGG |
| *ccl2* | TAAAAACCTGGATCGGAACCAAA | GCATTAGCTTCAGATTTACGGGT |
| *chil3* | CAGCATATGGGCATACCTTT | CAGACCTCAGTGGCTCCTT |
| *retnlg* | AGGAACTTCTTGCCAATCG | GCCTGAAGCCGTGATACT |
| *s100a9* | CAGCATCATACACTCCTCAAAG | AATGGTGGAAGCACAGTT |
| *itgam* | AAACCACAGTCCCGCAGAGA | CGTGTTCACCAGCTGGCTTA |
| *ubd* | CCAGATCCTTCTGCTAGACTCC | ACTCCACCAGAAACAAGGGCAG |
| *il6* | TAGTCCTTCCTACCCCAATTTCC | TTGGTCCTTAGCCACTCCTTC |
| *Rn18s* | GCAATTATTCCCCATGAACG | GGCCTCACTAAACCATCCAA |
| *Tnfa* | ACCCTCACACACTCAGATCATCTTC | TGGTGGTTTGCTACGACGT |
| *Nef* | ATGGGTGGCAAGTGGTCAA | TCAGCAGTTCTTGAAGTACTC |
| *Env* | TGTGTAAAATTAACCCCACTCTG | ACAACTTATCAACCTATAGCTGGT |

**List of primers used in the study.**
